# A GPCR signaling pathway in insect odor detection

**DOI:** 10.1101/2025.10.03.680299

**Authors:** Suguru Takagi, Liliane Abuin, Jérôme Mermet, Daehan Lee, Richard Benton

## Abstract

Odor detection differs fundamentally in vertebrates, which use G protein-coupled receptors (GPCRs)^1,2^, and insects, which employ ion channels^3,4^. Here, we report the first evidence for a GPCR defining tuning properties of insect olfactory sensory neurons. Single-cell transcriptomics of the *Drosophila melanogaster* antenna identified selective expression of the Gγ30A subunit in acid-sensing Ir64a-DC4 neurons^5,6^. Gγ30A is essential for broadening responses to long-chain acids, acting with Gα_s_, Gβ13F, adenylate cyclase Ac13E and the Cngl channel. We further discovered that Cirl, a latrophilin-family GPCR^7^, is broadly-transcribed in the antenna but the protein is localized only in Ir64a-DC4 sensory cilia, dependent upon Gγ30A, but not Ir64a. Importantly, loss of Cirl also narrows Ir64a-DC4 tuning properties. Homologous neurons in *Drosophila sechellia* naturally exhibit narrow acid tuning, despite functional conservation of Ir64a; these differences correlate instead with lower expression of metabotropic components. Our findings reveal unexpected roles for GPCR/metabotropic signaling in olfactory detection and divergence in insects.

## Introduction

A hallmark of most olfactory sensory neurons (OSNs) across vertebrates and insects is the selective expression of a single receptor type, which defines the neuronal odor response profile^8–10^. Neurons expressing a given receptor are also distinguished by their projection to a discrete glomerulus within the primary olfactory center (olfactory bulb in mammals; antennal lobe in insects)^11–13^. One intriguing exception is the population of OSNs in *Drosophila melanogaster* that express Ir64a, a member of the variant ionotropic glutamate receptor (iGluR) family of insect chemosensory receptors^14,15^. The sensory endings of these neurons are housed in porous cuticular sensilla in the sacculus^5,6,16^, a multichambered pocket within the antenna^17^ (**Figure 1a**). While these neurons all express Ir64a – together with the essential co-receptor Ir8a^5,16^ – they comprise two subtypes. These subtypes are housed in morphologically-distinct sensilla (Grooved Sensilla 1 and 2 (GS1 and GS2)) and project to discrete glomeruli, DC4 and DP1m (**Figure 1a**)^5,6,18^. Notably, the subtypes (hereafter, “Ir64a-DC4” and “Ir64a-DP1m” neurons) also exhibit different odor-response profiles: Ir64a-DC4 neurons are responsive mainly to acidic odors and Ir64a-DP1m neurons are more broadly tuned to diverse chemical classes^5,6^. Our study was motivated to understand the molecular basis underlying this differential odor tuning.

**Figure 1.**
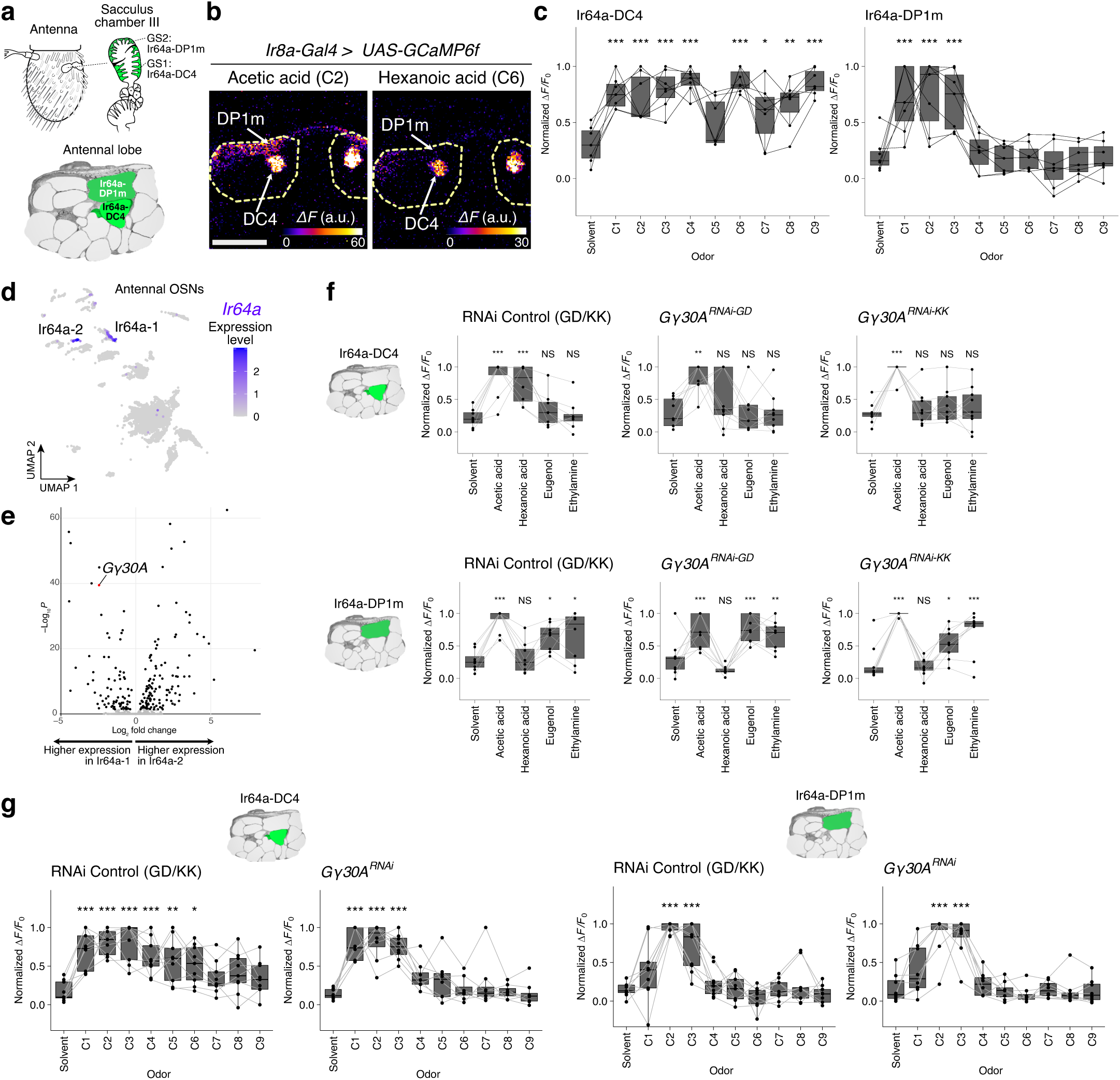
Differentially-expressed Gγ30A is required for acid-tuning divergence in Ir64a neuron subtypes. **a** Schematics of the peripheral and central anatomical organization of the two subtypes of Ir64a neurons. The sacculus cartoon was redrawn from^17^; GS1 and GS2 denote the distinct morphological classes of grooved sensilla housing the two Ir64a neuron subtypes. The antennal lobe model was generated from the dataset in^11,53^. **b** Representative odor-evoked calcium responses in the axon termini of Ir64a-DC4 and Ir64a-DP1m neurons. Dotted yellow line marks the outline of the antennal lobe. Scale bar = 50 µm. Genotype: *w;Ir8a-Gal4/UAS-GCaMP6f*. **c** Quantifications of calcium responses (normalized peak *ΔF/F_0_*, here and elsewhere) to a series of linear carboxylic acids (C1: formic, C2: acetic, C3: propionic, C4: butyric, C5: valeric, C6: hexanoic, C7: heptanoic, C8: octanoic, C9: nonanoic acids). *n* = 7 animals. Genotype as in (b). For these and all other box plots, the center line represents the median, the box bounds represent the first and third quartiles, and whiskers depict at maximum 1.5× the interquartile range; individual data points are overlaid. **d** UMAP of an antennal sensory neuron developmental snRNA-seq atlas (showing late developmental phase cells only), illustrating the expression of *Ir64a* in two cell clusters (expression levels have arbitrary units). Data from^19^. **e** Volcano plot illustrating differentially-expressed genes between the two subtypes of Ir64a neurons (pseudobulk analysis, *x* axis = log_2_FC(subtype 2/1), *y* axis = - log_10_(adjusted *P*), log_2_FC > 0, % of positive nuclei > 0.15). Genes significantly enriched in subtype 1 and 2 (adjusted *P* < 0.05) are highlighted in black. See also **Extended Data Table 1**. **f** Quantifications of calcium responses to the indicated odors upon knock-down of *Gγ30A*. *n* = 8 (control), 9 (*Gγ30A^RNAi-GD^*), 9 (*Gγ30A^RNAi-KK^*) animals. Genotypes: *w/w,UAS-Dcr-2;Ir64a-Gal4,UAS-GCaMP6f/+* (control) | *w/w,UAS-Dcr-2;Ir64a-Gal4,UAS-GCaMP6f/UAS-Gγ30A^RNAi-GD^*(*Gγ30A^RNAi-GD^*) | *w/w,UAS-Dcr-2;Ir64a-Gal4,UAS-GCaMP6f/UAS-Gγ30A^RNAi-KK^* (*Gγ30A^RNAi-KK^*). **g** Quantifications of calcium responses to the indicated odors upon *Gγ30A* knock-down. *n* = 9 (control) and 8 (*Gγ30A^RNAi^*) animals. Genotypes are the same as in (F) (control and *Gγ30A^RNAi-KK^*). Dunnett’s test (two-sided, control: solvent), \*\*\**P* < 0.001, \*\**P* < 0.01, \**P* < 0.05, otherwise *P* > 0.05.

## Results

### Distinct acid responses of Ir64a neuron subtypes require Gγ30A

We first re-examined the response profile of Ir64a neuron subtypes. As Ir64a neurons are inaccessible to peripheral electrophysiological recordings, we performed two-photon calcium imaging from OSN axon termini in the antennal lobe, using the *Ir8a*-Gal4 driver to express GCaMP6f in these neurons (**Extended Data** Figure 1a). Ir64a-DC4 neurons were previously described as broadly-tuned acid sensors^5,6^, and we confirmed robust responses to linear carboxylic acids of diverse chain-length, such as acetic acid (C2) and hexanoic acid (C6) (**Figure 1b-c**), as well as a few additional non-acid ligands (**Extended Data** Figure 1b). Ir64a-DP1m neurons also responded to acids, but this was restricted to short-chain acids, as carboxylic acids containing >3 carbons did not evoke significant responses (**Figure 1b-c**). Consistent with previous observations^5,6^, Ir64a-DP1m neurons also responded to many other chemicals (**Extended Data** Figure 1b). Thus, while the two populations of Ir64a neurons have some common ligands, their tuning profiles are different: Ir64a-DC4 neurons have broader tuning to acid odors, while Ir64a-DP1m neurons have broader tuning to other chemical classes of stimuli.

To understand the mechanism(s) underlying these distinct response profiles, we sought genes that are differentially expressed between the neuronal subtypes using an antennal single-nucleus RNA-sequencing (snRNA-seq) atlas^19^. Two Ir64a-expressing cell clusters (“1” and “2”) were annotated in this dataset^19^ (**Figure 1d**), which presumably correspond to the two Ir64a neuron subtypes. Beyond *Ir64a* and *Ir8a* (and the broadly-expressed *Ir25a*^16,20^) we did not detect transcripts for any additional known chemosensory receptors in either of the neuron types^19^. Out of 2034 detected genes, 77 were significantly more highly expressed in Ir64a-1 and 121 genes in Ir64a-2 cell clusters (**Figure 1e** and **Extended Data Table 1**). After excluding genes that are broadly-expressed in the antenna (which seemed unlikely to contribute to specific odor tuning properties) and those encoding proteins with irrelevant or indirect functions (e.g., transcription factors), we screened candidates by knocking down their expression through transgenic RNAi induced by the *Ir64a*-Gal4 driver. We examined neuronal responses to acetic and hexanoic acids (which distinguish Ir64a-DC4 and Ir64a-DP1m acid tuning breadth (**Figure 1b**)), as well as eugenol and ethylamine (which selectively activate Ir64a-DP1m neurons (**Extended Data** Figure 1b)). As a positive control we first verified that *Ir64a^RNAi^* abolished all tested odor responses in Ir64a-DC4 and Ir64a-DP1m neurons (**Extended Data** Figure 1c). In the course of screening, *Gγ30A^RNAi^* produced a notable phenotype: in Ir64a-DC4 neurons, hexanoic acid responses were abolished while acetic acid responses were maintained, while in Ir64a-DP1m neurons, responses to all stimuli were still observed (**Figure 1f**). As *Gγ30A* is enriched in Ir64a-1 (**Figure 1e**), we infer that this cell type likely corresponds to the Ir64a-DC4 neurons.

We extended our investigation of the role of Gγ30A in acid sensing by testing a broader range of organic acids from formic acid (C1) to nonanoic acid (C9). *Gγ30A^RNAi^* resulted in restriction of sensitivity of Ir64a-DC4 to short-chain acids (<C4), similar to the acid tuning profile of wild-type Ir64a-DP1m (**Figure 1g**). These results suggest that Gγ30A broadens the tuning of Ir64a-DC4 neurons to encompass longer acids. Other, as-yet unidentified, factors presumably determine the broader sensitivity of Ir64a-DP1m neurons to diverse chemicals (**Extended Data** Figure 1b); these might exist among the other differentially-expressed genes in Ir64a populations (**Figure 1e**), although we cannot exclude that tuning properties are influenced by differences in associated non-neuronal cells^21^.

### A G**α**_s_ signaling pathway is required for long-chain acid responses in Ir64a-DC4 neurons

*Gγ30A* encodes a γ subunit of heterotrimeric G proteins^22,23^, leading us to ask with which G protein α and β subunits it might act. Using the antennal snRNA-seq dataset^19^, we first examined the expression of all *D. melanogaster* G protein subunits genes in Ir64a neurons, detecting transcripts for four of six *α* subunits (*Gα_s_*, *Gα_o_*, *Gα_q_*, *cta*), one of three β subunits (*Gβ13F*); in addition to *Gγ30A* we also detected the other γ subunit (*Gγ1*) (**Figure 2a and Extended Data** Figure 2a).

**Figure 2.**
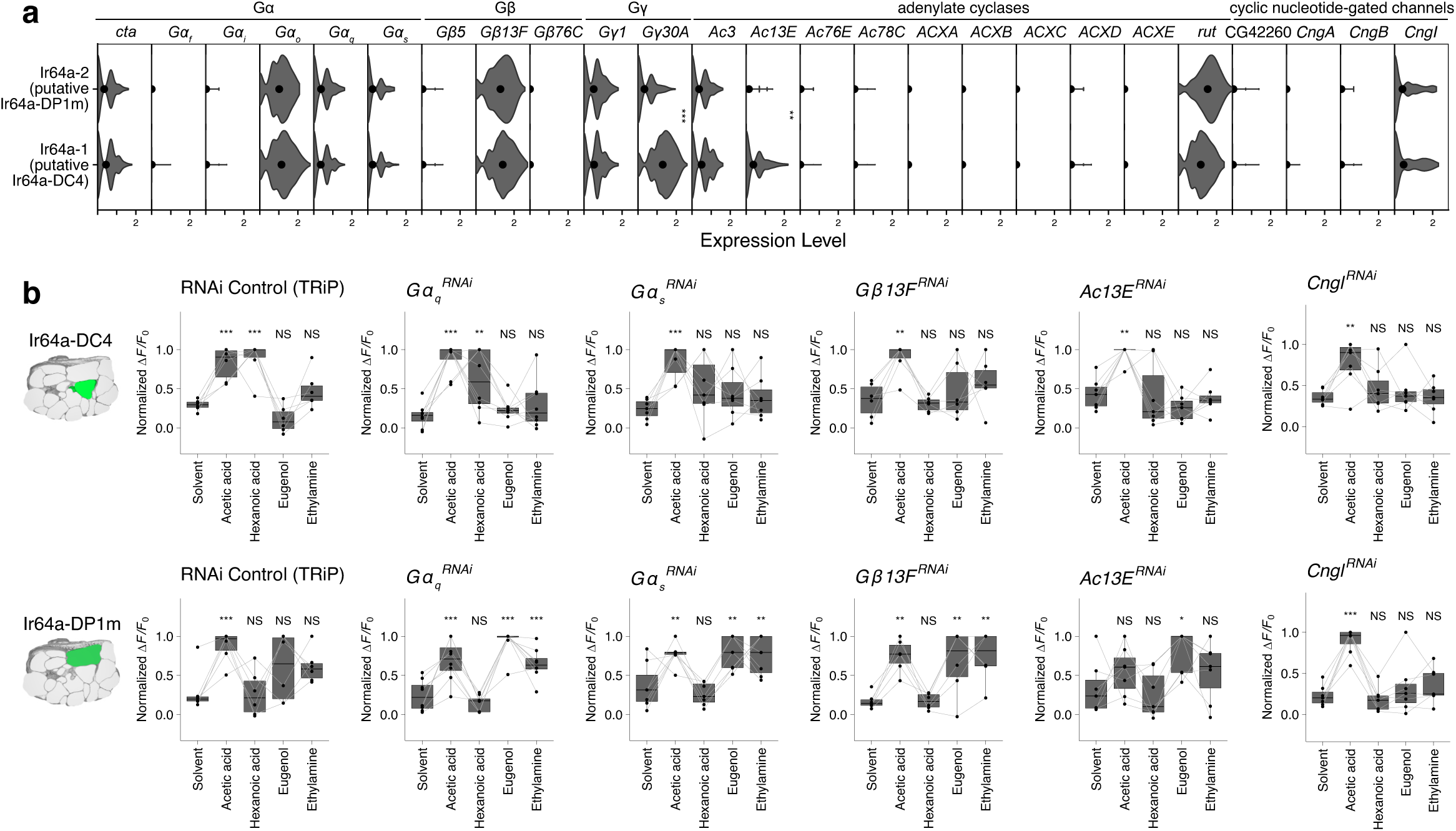
A metabotropic pathway required for acid-tuning divergence in Ir64a neuron subtypes. **a** Violin plot showing the expression of G protein subunits, adenylate cyclases, and cyclic nucleotide-gated ion channel protein transcripts in the two Ir64a neuron subtypes. Dots show the mean expression (arbitrary units) for each gene and neuron type, here and elsewhere. **b** Quantifications of calcium responses to the indicated odors upon knock-down of G protein subunits and adenylate cyclase genes. *n* = 6 (control), 8 (*Gα_q_^RNAi^*), 7 (*Gα_s_^RNAi^*), 6 (*Gβ13F^RNAi^*), 7 (*Ac13E^RNAi^*), 7 (*Cngl^RNAi^*) animals. Genotypes: *y,sc,v,sev/w,UAS-Dcr-2;Ir64a-Gal4,UAS-GCaMP6f/+;UAS-mCherry.VALIUM10/+* (control) | *y,v/w,UAS-Dcr-2;Ir64a-Gal4,UAS-GCaMP6f/+;UAS-Gα_q_^RNAi^/+* (*Gα_q_^RNAi^*) *y,v/w,UAS-Dcr-2;Ir64a-Gal4,UAS-GCaMP6f/+;UAS-Gα_s_^RNAi^/+* (*Gα_s_^RNAi^*) | *y,sc,v,sev/w,UAS-Dcr-2;Ir64a-Gal4,UAS-GCaMP6f/+;UAS-Gβ13F^RNAi^/+* (*Gβ13F^RNAi^*) | *y,sc,v,sev/w,UAS-Dcr-2;Ir64a-Gal4,UAS-GCaMP6f/UAS-Ac13E^RNAi^*(*Ac13E^RNAi^*) | *y,v/w,UAS-Dcr-2;Ir64a-Gal4,UAS-GCaMP6f/+;UAS-Cngl^RNAi-TRiP^/+* (*Cngl^RNAi^*). Dunnett’s test (two-sided, control: solvent), \*\*\**P* < 0.001, \*\**P* < 0.01, \**P* < 0.05, otherwise *P* > 0.05.

We tested the role of all expressed Gα and Gβ subunits by RNAi in Ir64a neurons (**Extended Data** Figure 2b). Similar to loss of *Gγ30A*, *Gα_s_^RNAi^* (but not *Gα_q_^RNAi^*) and *Gβ13F^RNAi^* selectively diminished hexanoic acid responses in Ir64a-DC4 neurons, while not affecting responses to the test odors in Ir64a-DP1m neurons (**Figure 2b**). These data imply that Gγ30A forms a complex with G*α*_s_ and Gβ13F to influence the odor tuning properties of Ir64a-DC4 neurons. We note, however, that neither *Gα_s_* nor *Gβ13F* are differentially expressed between the two Ir64a neuron subtypes (**Figure 2a** and **Extended Data Table 1**), suggesting that the enrichment of *Gγ30A* in Ir64a-DC4 neurons underlies the divergent acid responses. Surveying G protein subunit expression across all populations of antennal sensory neurons, revealed that *Gγ30A* transcription displays remarkable heterogeneity compared to other G protein subunit genes (**Extended Data** Figure 2a), raising the possibility that it has a role in the engagement of heteromeric G proteins in other neuronal populations.

The involvement of Gα_s_ implies the downstream activation of adenylate cyclase to produce the second messenger cAMP^24,25^. We tested this hypothesis by RNAi of the three adenylate cyclase genes (*Ac3*, *Ac13E*, *rut*) expressed in Ir64a OSNs (**Figure 2a, Extended Data** Figure 2a-b). *Ac13E^RNAi^* selectively abolished hexanoic acid responses in Ir64a-DC4 neurons (**Figure 2b**), similar to knock-down of G protein subunits. *Ac13E* exhibits higher expression in the inferred Ir64a-DC4 population, similar to (albeit less pronounced than) the differential expression of *Gγ30A* (**Figure 2a** and **Extended Data Table 1**). Unlike the G protein subunits, however, Ac13E appears to have a role in Ir64a-DP1m, as *Ac13E^RNAi^* led to milder, albeit somewhat variable, effects on some odor responses in these neurons (**Figure 2b**).

Given that a heterotrimeric G protein and an adenylate cyclase contribute to odor-evoked neuronal activity in Ir64a-DC4 neurons, we hypothesized that this pathway would ultimately lead to neuronal depolarization by acting on a cyclic nucleotide-gated channel. Of the four such channels in *D. melanogaster*, only one, *Cngl*, is expressed in Ir64a neurons (**Figure 2a** and **Extended Data** Figure 2a). Consistent with our hypothesis, *Cngl^RNAi^* abolished responses to hexanoic acid, but not acetic acid, of Ir64a-DC4 neurons (**Figure 2b**). However, loss of this channel also led to diminished responses of Ir64a-DC4 neurons to formic and propionic acids, and loss of responses of Ir64a-DP1m neurons to eugenol and ethylamine (**Figure 2b and Extended Data** Figure 2c), perhaps reflecting additional roles of this channel in neuronal signaling. Nevertheless, taken together, these results suggest that Ir64a-DC4 neurons require a G protein- and cAMP-dependent signaling cascade to be able to respond to long-chain carboxylic acids.

### The GPCR Cirl is selectively expressed in sensory cilia of Ir64a-DC4 neurons

The involvement of G protein signaling in odor tuning of Ir64a-DC4 neurons begged the question of how such a metabotropic pathway might be activated. There is limited evidence for vertebrate iGluRs coupling to G proteins^26,27^, and no obvious G protein interaction domains in Irs, leading us to pursue the hypothesis that a canonical GPCR functions upstream of G protein signaling – and in concert with Ir64a/Ir8a – to define neuronal response properties.

Of the 117 genes encoding predicted GPCRs in *D. melanogaster*, we found only 16 were expressed at more than trace levels (mean expression >0.3) in Ir64a-DC4 OSNs (**Figure 3a, Extended Data** Figure 3a). Most of these GPCRs have well-established roles in neurotransmission and axon guidance (e.g., Dop1R1, fz, fz2, Oct-TyrR, stan), or as neuropeptide receptors (e.g., Dh31-R, SIFaR, SPR). Our attention was caught by *Calcium-independent receptor for α-latrotoxin* (*Cirl*)^7^, one of the most highly-expressed GPCR genes in Ir64a-DC4 neurons, which encodes an adhesion GPCR of the latrophilin family^28^. In *D. melanogaster* larvae, Cirl functions in the peripheral nervous system in modulating electrophysiological responses of mechanosensory neurons^7,29^, raising the possibility of an analogous function of this GPCR in OSNs.

**Figure 3.**
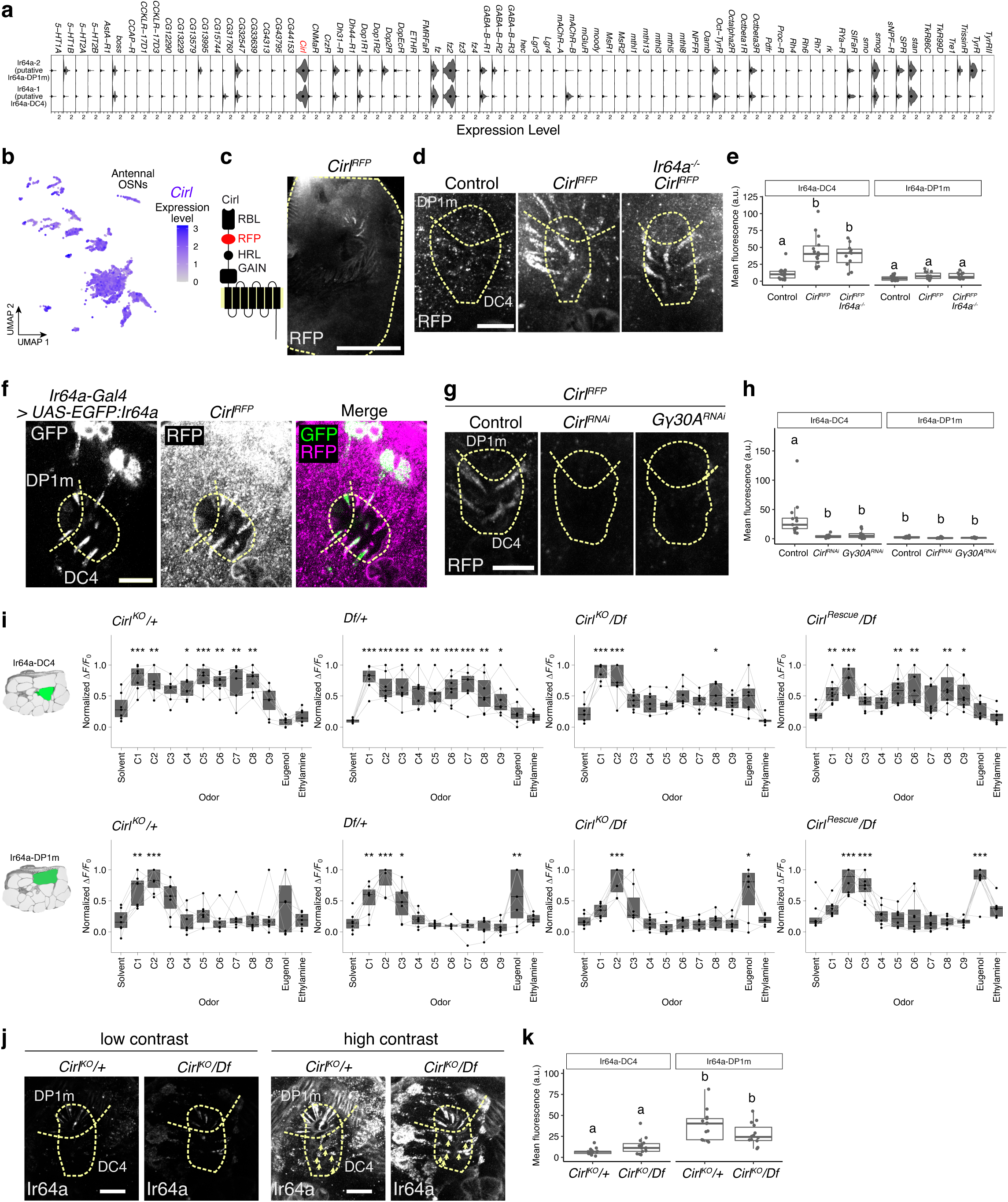
Sensory cilia localization and function of Cirl/Latrophilin in Ir64a-DC4 neurons. **a** Violin plot showing the expression of all GPCR genes detected (at even trace levels; normalized data with regression fit) in the two Ir64a neuron subtypes. *Cirl* is highlighted in red. **b** UMAP (as in Figure 1d) showing the broad expression of *Cirl* in all OSN clusters. **c** Left: schematic of Cirl protein structure, indicating the position of the RFP tag. Right: representative image of the localization of Cirl^RFP^ fusion protein (*w^11^*^18^;*Cirl^RFP.V5^*) in a whole-mount antenna, visualized with anti-RFP. Scale bar = 50 µm. **d** Representative images of the localization of Cirl^RFP^ (visualized with anti-RFP) in sacculus chamber III in the presence or absence of Ir64a. Dotted line outlines the sacculus chamber III. Scale bar = 10 µm. Genotypes: *w^1118^* (control) *| w^1118^*_;_*Cirl^RFP.V5^ | w^1118^*_;_*Cirl^RFP.V5^*_;*I*_*r64a^MB05283^*. **e** Quantification of mean fluorescent signal of Cirl^RFP^ from the two sensillar types (GS1 and GS2) in sacculus chamber III, corresponding to Ir64a-DC4 and Ir64a-DP1m dendrites^5,6^. *n* = 15 (*w^1118^*(control)), 14 (*w^1118^*;*Cirl^RFP.V5^*), 11 (*w^1118^*;*Cirl^RFP.V5^*;*Ir64a^MB05283^*) antennae. **f** Co-localization of EGFP-tagged Ir64a and Cirl^RFP^ (visualized with anti-GFP and anti-RFP) in sacculus chamber III in the antenna. While GFP signal can be readily detected in soma and sensory dendrites, the RFP signal is mostly evident only in the sensory endings. Scale bar = 10 µm. Genotype: *w;Cirl^RFP.V5^/UAS-SS:Ir64a:EGFP;Ir64a-Gal4/+*. Some Cirl-positive sacculus sensilla lack GFP signal, likely due to incomplete expression of the *Ir64a-Gal4* driver. **g** Left: Representative images of Cirl^RFP^ localization upon *Cirl* and *Gγ30A* knock-down in Ir64a neurons, visualized with anti-RFP. Scale bar = 10 µm. Genotypes: *w/w,UAS-Dcr-2;Cirl^RFP.V5^/+;Ir64a-Gal4/+* (control); *w/w,UAS-Dcr-2;Cirl^RFP.V5^/UAS-Cirl^RNAi-GD^;Ir64a-Gal4/+* (*Cirl^RNAi^*) | *w/w,UAS-Dcr-2;Cirl^RFP.V5^/UAS-Gγ30A^RNAi-GD^;Ir64a-Gal4/+* (*Gγ30A^RNAi^*). **h** Quantification of mean fluorescent signal from each sensilla type. *n* = 13 (control), 12 (*Cirl^RNAi^*), 13 (*Gγ30A^RNAi^*) antennae. Genotypes as in (g). **i** Quantifications of calcium responses to the indicated odors in *Cirl* null mutant animals. *n* = 7 animals. Genotypes: *w;UAS-GCaMP6f,Cirl^KO^/+;Ir64a-Gal4/+* (*Cirl^KO^/+*) | *w;UAS-GCaMP6f/Df(2R)Exel8047;Ir64a-Gal4/+* (*Df(2R)Exel8047/+*) | *w;UAS-GCaMP6f,Cirl^KO^/Df(2R)Exel8047;Ir64a-Gal4/+* (*Cirl^KO^/Df(2R)Exel8047*) | *w;UAS-GCaMP6f,Cirl^Rescue^/Df(2R)Exel8047;Ir64a-Gal4/+* (*Cirl^Rescue^/Df(2R)Exel8047*) For (E) and (H), Tukey’s HSD test, different letters indicate statistically significant differences with *P_adj_*<0.0001. For (I), Dunnett’s test (two-sided, control: solvent), \*\*\**P* < 0.001, \*\**P* < 0.01, \**P* < 0.05, otherwise *P* > 0.05. **j** Left: Representative images of Ir64a expression (visualized with anti-Ir64a) in the sacculus chamber III sensilla in the antennae of *Cirl* heterozygous (*Cirl^KO^/+*) and hemizygous (*Cirl^KO^/Df*) mutants. As signal intensity differed considerably between Ir64a-DC4 and Ir64a-DP1m cilia, both low and high contrast images are shown. Dotted lines outline the sacculus chamber III. Arrowheads point to Ir64a signals in Ir64a-DC4 cilia. Scale bar = 10 µm. **k** Quantification of mean fluorescent signal from each sensilla type. *n* = 13 (*Cirl^KO^/+*) and 14 (*Cirl^KO^/Df(2R)Exel8047*) antennae. Tukey’s HSD test, different letters indicate statistically significant differences with *P_adj_*<0.05.

*Cirl* transcripts are broadly detected in antennal sensory neurons (**Figure 3b**). Consistently, a transcriptional reporter for this gene, *Cirl^Gal^*^4^, is expressed in many different types of neurons in the antenna (**Extended Data** Figure 3b). No Cirl antibodies are available^30^, so to detect Cirl protein, we used a reporter line, *Cirl^RFP.V5^*^29,31^, under the control of the endogenous promoter, in which *RFP* is inserted between regions encoding two of the extracellular domains in the seven-transmembrane domain Cirl isoform (**Figure 3c**), which avoids disrupting localization or function as well as over-expression artefacts^29–32^. Strikingly, Cirl^RFP.V5^ was detected only in neurons located close to sacculus chamber III (**Figure 3c**). High-resolution imaging revealed the presence Cirl in the sensory dendrites housed in GS1 sensilla (see Methods), corresponding to those housing Ir64a-DC4 neurons^5^ (**Figure 3d-e**). This signal overlapped with that of EGFP-tagged Ir64a expressed transgenically in Ir64a neurons, suggesting that Cirl expression is derived from these neurons (**Figure 3f**). Consistent with this possibility, transgenic RNAi of *Cirl* under the control of *Ir64a*-Gal4 abolished Cirl protein signal (**Figure 3g-h**), confirming that the observed sensillar signal is derived from Ir64a neurons. In contrast to the specific protein signal in Ir64a sensory cilia in the antenna, Cirl protein was detected broadly in the central brain, including signal in the antennal lobe (**Extended Data** Figure 3c), matching the widespread transcription of this gene^33^.

To better understand why Cirl protein is only robustly detected in sensory cilia of Ir64a-DC4 neurons, we first examined Cirl^RFP.V5^ localization in *Ir64a* mutants (**Figure 3d-e**). Cirl^RFP.V5^ signal was still apparent in Ir64a-DC4 cilia in this mutant background, indicating that it localizes to this sensory compartment independently of this Ir. Given the differential expression of Gγ30A between Ir64a neuron subtypes (**Figure 2a**), we next tested whether the G protein subunit was required for Cirl localization. Indeed, *Gγ30A^RNAi^* resulted in loss of Cirl^RFP.V5^ signal (**Figure 3g-h**), implying that Gγ30A is required, directly or indirectly, in the localization of Cirl to the sensory endings of Ir64a-DC4 neurons.

Together, these results highlight a remarkable post-transcriptional regulation that restricts the presence of the GPCR Cirl to the sensory cilia of only a small subset of the large number of neurons that transcribe the corresponding gene.

### Cirl is required for long-chain acid responses in Ir64a-DC4 neurons

The selective protein expression of Cirl strongly pointed to the candidacy of this GPCR in acting with Ir64a in olfactory responses of Ir64a-DC4 neurons. To test this hypothesis, we performed calcium imaging in Ir64a neurons in *Cirl* null mutant animals. Importantly, we found that in the absence of Cirl, Ir64a-DC4 neurons lose the broad acid tuning observed in control animals (**Figure 3i**), similar to the knock-down of various metabotropic signalling components (**Figure 2b**). The broad acid tuning was largely restored by genetic rescue of *Cirl* (**Figure 3i**).

*Cirl* transcripts are detected throughout development, contrasting with *Ir64a*, whose transcription initiates only at late developmental stages (**Extended Data** Figure 3d). Cirl has been implicated in synapse assembly^34^ – most similar to its best-described roles in vertebrates^28^ – and in planar cell polarization during development^32^. To exclude a possible developmental or synaptic origin of the *Cirl* mutant phenotype, we first examined Ir64a neuron morphology in the antennal lobe in *Cirl* mutants. No obvious differences compared to controls were observed (**Extended Data** Figure 3e), indicating that the specification and wiring of these neurons are not markedly affected by the loss of this GPCR. No gross anatomical changes were detected elsewhere in the antennal lobe. We next induced *Cirl* loss-of-function in late developmental stages by RNAi using the *Ir64a-Gal4* driver. Similar to the *Cirl* mutant, *Cirl^RNAi^* diminished the hexanoic acid responses in Ir64a-DC4 neurons (**Extended Data** Figure 3f), consistent with a cell-autonomous role for Cirl in mature Ir64a neurons. Finally, we examined Ir64a localization in *Cirl* mutants. Ir64a signal in Ir64a-DC4 neurons was extremely weak, even in controls, and this was unchanged (or even slightly, but non-significantly, elevated) in *Cirl* mutants (**Figure 3j-k**). This result indicates that Cirl is not essential for sensory cilia formation of these neurons nor for targeting of Ir64a to this cellular compartment.

### Divergence in *Drosophila sechellia* Ir64a-DC4 OSN acid responses through an Ir64a-independent mechanism

Ir64a-DC4 neurons are required for behavioral avoidance of acids by *D. melanogaster*^6^. By contrast, in the close relative *Drosophila sechellia*, hexanoic acid is attractive^35^, reflecting the abundance of this stimulus in this species’ host noni fruit^36,37^. This attraction is mediated in part by the evolution of novel sensitivity of *D. sechellia* Ir75b to hexanoic acid (compared to *D. melanogaster* Ir75b, which responds only to shorter-chain acids) through amino-acid changes in its ligand-binding domain^35,38^.

We wondered whether Ir64a-DC4 neurons might also exhibit differences in response properties in *D. sechellia* as part of its behavioral adaptation. We generated an *Ir8a-*Gal4 driver in *D. sechellia* and used this to express GCaMP6f, enabling us to record odor-evoked activity in Ir64a-DC4 and Ir64a-DP1m in this species (**Figure 4a**). Acetic acid strongly activated both Ir64a-DC4 and Ir64a-DP1m neurons in *D. sechellia,* similar to *D. melanogaster* (**Figure 4a-b**). However, *D. sechellia* Ir64a-DC4 neurons did not respond to hexanoic acid, unlike the homologous neurons in *D. melanogaster* and instead more similar to Ir64a-DP1m neurons (**Figure 4a-b**). Indeed, quantitative comparison of the response profile of Ir64a-DC4 and Ir64a-DP1m neurons for a panel of odors (**Extended Data** Figure 1d) revealed much greater similarity in *D. sechellia* compared to *D. melanogaster* (**Figure 4c, Extended Data** Figure 1b**, Extended Data** Figure 1d**, and Extended Data** Figure 4a).

**Figure 4.**
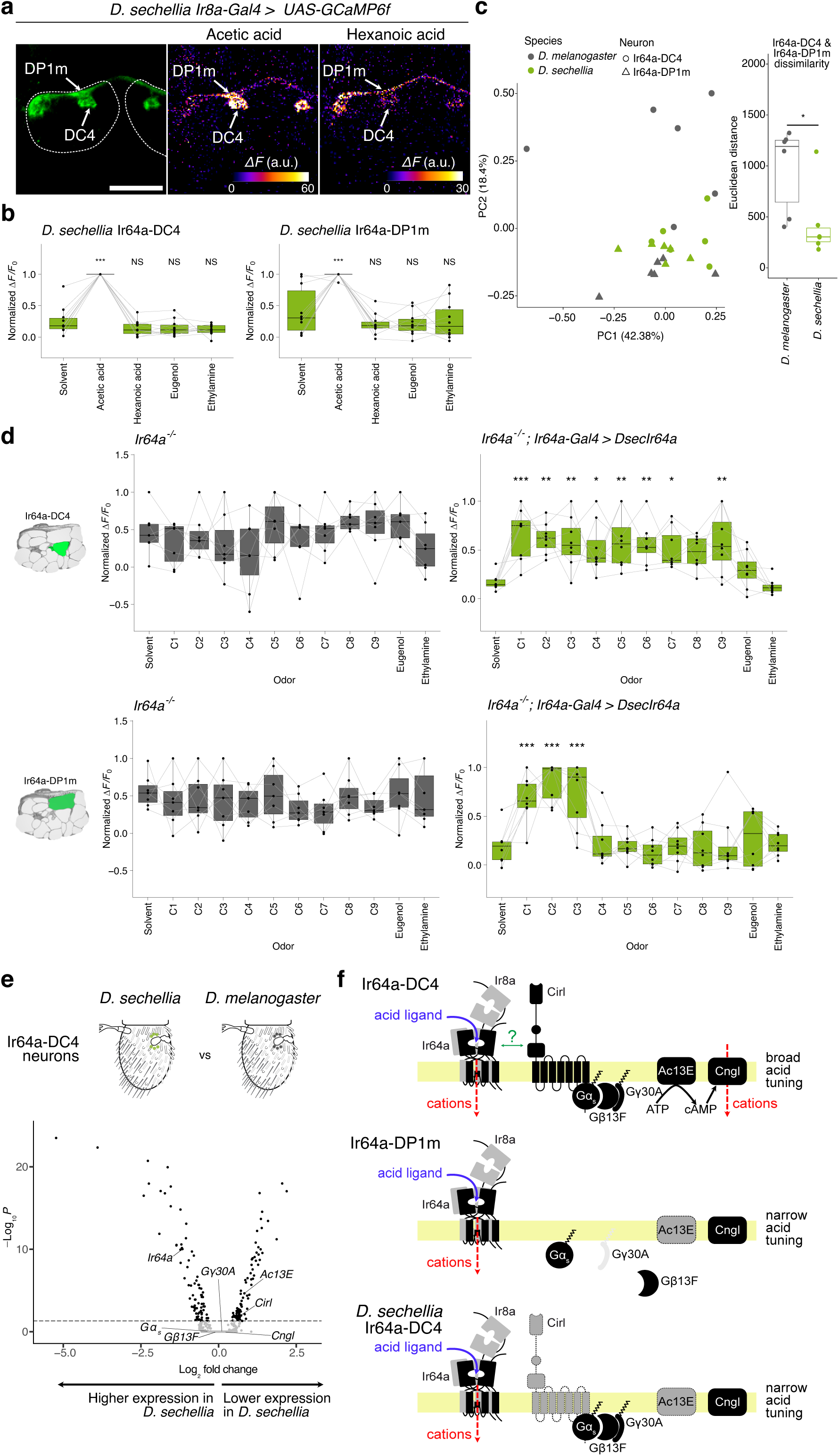
Evolutionary divergence of acid-tuning of Ir64a neurons. **a** Top left: Expression of *UAS-GCaMP6f* under the control of *Ir8a-Gal4* in *D. sechellia*. Arrows indicate the two Ir64a glomeruli. Top right: representative odor-evoked calcium responses in the axon termini of Ir64a-DC4 and Ir64a-DP1m neurons in *D. sechellia*. Genotype: *w/w*;*UAS-GCaMP6f;Ir8a-Gal4/+* (chromosome unidentified). As observed previously^35^, DP1m is smaller in *D. sechellia* than *D. melanogaster*, which might reflect a lower number of neurons innervating this glomerulus. **b** Quantifications of calcium responses to the indicated odors. *n* = 10 animals. Genotype: *w/w,UAS-GCaMP6f;Ir8a-Gal4/+* (chromosome unidentified). **c** Left: Principal component analysis of odor tuning profile based on broader odor responses (**Extended Data** Figure 1b,d) in both Ir64a subtypes (DC4 and DP1m) and both species (*D. melanogaster* and *D. sechellia*). Right: Quantifications of the Euclidean distance between Ir64a-DC4 and Ir64a-DP1m neurons in each species. *n* = 6 animals. **d** Quantifications of calcium responses to the indicated odors in animals expressing *D. sechellia Ir64a* allele (*DsecIr64a*) in *D. melanogaster* Ir64a neurons lacking endogenous receptor expression. *n* = 7 (*Ir64a^-/-^*) and 8 (*Ir64a^-/-^, Ir64a-Gal4>DsecIr64a*) animals. Genotypes: *w;UAS-GCaMP6f/Ir64a-Gal4;Ir64a^MB05283^*(*Ir64a^-/-^*) | *w;UAS-GCaMP6f/Ir64a-Gal4;UAS-DsecIr64a,Ir64a^MB05283^/Ir64a^MB05283^* (*Ir64a^-/-^,Ir64a-Gal4>DsecIr64a*). **e** Volcano plot illustrating differentially-expressed genes between *D. melanogaster* and *D. sechellia* in Ir64a neuron subtype 1 (presumed Ir64a-DC4 neurons). Genes significantly enriched in *D. melanogaster* or *D. sechellia* are highlighted in black. See also **Extended Data Table 2**. **f** Model of the Ir64a neuron metabotropic pathway function in different Ir64a neuron subtypes and species. Lower expression of *Ac13E* and *Cirl*, and of *Gγ30A*, is illustrated with dark and pale grey shading, respectively.

A plausible reason underlying interspecific tuning differences of Ir64a-DC4 neurons is that the Ir64a receptor is functionally distinct between *D. melanogaster* and *D. sechellia*, as for Ir75b^35^. To test this possibility, we expressed *D. sechellia Ir64a* in *D. melanogaster* Ir64a OSNs lacking the endogenous receptor. As expected, both subtypes of *Ir64a* mutant neurons lack all odor responses (**Figure 4d**). *D. sechellia* Ir64a conferred sensitivity to a broad range of acidic odors when expressed in these *D. melanogaster* Ir64a-DC4 OSNs (**Figure 4d**), in contrast to the native tuning profile of *D. sechellia* Ir64a-DC4 neurons. By contrast, in *D. melanogaster* Ir64a-DP1m OSNs, *D. sechellia* Ir64a only conferred responses to short-chain aids (**Figure 4d**). These observations imply that the tuning differences of Ir64a-DC4 neurons in *D. melanogaster* and *D. sechellia* cannot be explained by functional differences in Ir64a.

To investigate other potential explanations for the interspecific differences in Ir64a tuning we sought to compare the transcriptional profiles of the Ir64a-DC4 neurons. We used snRNA-seq to generate antennal atlases of adult *D. sechellia* and *D. melanogaster*. Within the annotated OSN types (see Methods), we identified two subtypes of Ir64a neurons in both species, as observed in the previous developmental atlas^19^. Comparison of the transcriptomes of the presumed Ir64a-DC4 neurons identified 166 differentially-expressed genes between species (**Figure 4e and Extended Data Table 2**). *Ir64a* expression was higher in *D. sechellia* compared to *D. melanogaster* (**Figure 4e and Extended Data** Figure 4b), a finding corroborated by analysis of *Ir64a* expression in cross-species, bulk antennal RNA-seq datasets^39,40^. This observation indicates that lack of hexanoic acid responses in *D. sechellia* Ir64a-DC4 neurons also cannot be ascribed to lower receptor gene expression.

Given the similarity of acid responses of *D. sechellia* Ir64a-DC4 and Ir64a-DP1m neurons (**Figure 4c and Extended Data** Figure 4a), we hypothesized the existence of changes in the metabotropic pathway that is required in *D. melanogaster* for broadening the acid-tuning profile of Ir64a-DC4 neurons. None of the genes encoding G protein subunits (*Gγ30A*, *Gα_s_* and *Gβ13F*) or the cyclic nucleotide-gated channel (*Cngl*) were differentially expressed between species in these neurons (**Figure 4e and Extended Data** Figure 4b). However, *Ac13E* and *Cirl* are expressed at significantly lower levels in *D. sechellia* compared to *D. melanogaster* (**Figure 4e**); for both genes, this decrease reflects generally lower expression across antennal neuron types (**Extended Data** Figure 4b). It is possible that decreased *Ac13E* expression in *D. sechellia* explains some of the tuning differences observed in Ir64a-DP1m neurons between species (**Figure 4b and Figure 1d**).

Together, these observations raise the possibility that species differences in acid-tuning Ir64a-DC4 neurons are due not to changes in Ir64a function or expression, but rather to differential activity of the Cirl-dependent metabotropic pathway.

## Discussion

We have discovered a novel role for a GPCR, Cirl, and a metabotropic cascade in contributing to the odor-tuning properties of an insect OSN (**Figure 4f**). How might Cirl broaden the tuning properties of Ir64a-DC4 neurons to longer-chain acids? It seems unlikely to be acting as a receptor itself, as we and others^5,6^ have found that all sensory responses of these neurons depend upon Ir64a. We favor a model in which Cirl amplifies weak Ir64a-dependent responses evoked by longer-chain acid ligands through stimulation of a cAMP-dependent metabotropic pathway that ultimately gates Cngl. This amplification role is consistent with previous observations that high concentrations of longer-chain acids can evoke responses in Ir64a-DP1m neurons^6^. Mechanistically, such amplification might, in part, be through direct interaction of Cirl with Ir64a/Ir8a (**Figure 4f**), to modify odor-evoked conformational changes in this ionotropic complex. However, our implication of a metabotropic cascade downstream of Cirl points more concretely to a parallel pathway that increases basal neuronal depolarization, which could sensitize neurons to odor-evoked depolarization through Ir64a/Ir8a (**Figure 4f**). Testing such hypotheses will require functional reconstitution of these signaling pathways in cell types amenable to biochemical and electrophysiological analyses, which are difficult, if not impossible, in the endogenous neurons in the sacculus. Regardless of the precise mechanism, recent evidence that Cirl is involved in amplifying responses of mechanosensory neurons^7,31^ raises the possibility that Cirl has a general role in shaping sensory responsiveness across diverse modalities.

One striking observation in our work is the discordance between the broad transcription of *Cirl* and the selective detection of protein in Ir64a-DC4 neuron sensory cilia. The widespread neuronal expression of Cirl might reflect other, more general, functions of this protein at synapses, perhaps similar to mammalian homologs^28^. By contrast, the selective targeting of Cirl to the Ir64a-DC4 neuronal cilia appears to be important to engage a metabotropic pathway to define the broad acid tuning of these neurons compared to Ir64a-DP1m neurons. This targeting depends, at least in part, upon the selectively-expressed Gγ30A, highlighting a novel role for a Gγ subunit in GPCR localization and/or stabilization in sensory dendrites. However, we suspect that other factors are involved because Gγ30A is expressed in other OSN types where Cirl protein is not detected in the cilia endings (**Extended Data** Figure 2a). More generally, our findings provide an important demonstration that transcriptional properties of a gene do not necessarily predict the cellular specificity of function of the encoded protein.

In addition to the distinct tuning of Ir64a-DC4 and Ir64a-DP1m neurons within *D. melanogaster*, we have observed divergence of Ir64a-DC4 tuning properties between drosophilid species. Until now, evolutionary divergence in sensory neuron tuning properties has been exclusively associated with amino acid changes in the ligand-binding domains of the expressed odorant receptors (e.g.,^35,41,42^). The differential tuning of Ir64a-DC4 neurons in *D. melanogaster* and *D. sechellia* defines a novel case, as it cannot be ascribed to functional (or expression) differences of Ir64a orthologs. Rather, the narrower acid tuning of *D. sechellia* neurons might be associated with the lower expression of components of the Cirl-dependent metabotropic pathway, analogous to the mechanism underlying the acid tuning differences of Ir64a-DC4 and Ir64a-DP1m neurons. Interestingly, one transcriptional difference, of *Ac13E*, is observed both between Ir64a neuron subtypes within species and between homologous Ir64a-DC4 neurons across species, while other differentially-expressed genes are distinct, i.e., lower *Gγ30A* transcription in Ir64a-DP1m but lower *Cirl* in *D. sechellia* (**Figure 4f**).

Finally, our work adds an important advance to help clarify the long-term debate about the contribution of metabotropic signaling to insect olfactory transduction. It is widely accepted that insect chemosensory receptors – encompassing Irs and members of the insect Odorant/Gustatory receptor (Or/Gr) families – function as ligand-gated ion channels^3,4,43^. However, over the past decades, a number of G protein subunits and other metabotropic pathway components have been implicated genetically and/or pharmacologically in modulating the strength and/or temporal dynamics of chemosensory responses in various insect species^44–50^. How such signaling proteins act with ionotropic sensory receptors has remained unclear. Our discovery of Cirl as a GPCR that contributes to the tuning of an OSN population offers a new perspective in how metabotropic signaling modulates ionotropic chemosensory pathways. Beyond Ir64a neurons, we have revealed a diversity of expression patterns of metabotropic signaling components in antennal sensory neurons, from broad expression to remarkably heterogeneous expression (e.g., *Gγ30A* and *Cngl*) (**Extended Data** Figure 2a). These observations are suggestive of an unappreciated diversity of neuron-specific signaling pathways. An intriguing open question is whether additional GPCRs (**Extended Data** Figure 3a) have peripheral functions in other classes of insect chemosensory neurons.

## Acknowledgements

We are grateful to the Bloomington *Drosophila* Stock Center (NIH P40OD018537) and the Vienna *Drosophila* Resource Center for *D. melanogaster* stocks, and Greg Suh for Ir64a antibodies. We thank Bo-Yun Lee for bioinformatic support on single-cell analysis, and Nicole Scholz and Tobias Langenhan for advice. We thank Karen Menuz and members of the Benton laboratory for discussions and comments on the manuscript. S.T. is supported by Marie Skłodowska-Curie Actions Individual Fellowship (836783), an EMBO Long-Term Fellowship (ALTF 454-2019), and a Japanese Society for the Promotion of Science Overseas Research Fellowship (202360258). D.L. is supported by the National Research Foundation of Korea (NRF) grants funded by the Ministry of Science and ICT under Project Number RS-2023-00211007. Research in R.B.’s laboratory is supported by an ERC Advanced Grant (833548) and the Swiss National Science Foundation (310030_219185).

## Author contributions

S.T. and R.B. conceived the project. S.T. performed and analyzed all calcium imaging experiments, generated transgenic flies, and performed and analyzed histological experiments. L.A. performed molecular biology and histological experiments. J.M. analyzed the developmental snRNA-seq dataset in *D. melanogaster*. D.L. generated and analyzed the comparative antennal atlases of *D. melanogaster* and *D. sechellia*. R.B. supervised the project. S.T. and R.B. wrote the paper, with input from other co-authors.

## Methods

### Drosophila culture

*Drosophila* stocks were maintained on standard wheat flour/yeast/fruit juice medium under a 12 h light:12 h dark cycle at 25°C. For, *D. sechellia* strains, a few g of Formula 4-24® Instant Drosophila Medium, Blue (Carolina Biological Supply Company) soaked in noni juice (Raab Vitalfood or Tahiti Trader) were added on top of the standard medium. Strains used are listed in **Extended Data Table 3**.

### Transgenic animals

*Ir64a-Gal4 #183.6:* this is a third chromosome insertion of a previously-described construct^6^.

*D. sechellia Ir8a-Gal4*: a previously-described *D. melanogaster Ir8a promoter-Gal4* construct^16^ was injected into *D. sechellia w*^51^ embryos by WellGenetics. Transformants were screened based upon the *w^+^* marker.

*UAS-DsecIr64a*: the *D. sechellia Ir64a* ORF (*pEX-A258-NotI-Kozak-DsecIr64a-XbaI*) was synthesized by Eurofins Genomics. *NotI-Kozak-DsecIr64a-NheI* was amplified by PCR and digested with *NotI and NheI*. The fragment was inserted into *pJFRC7-20XUAS-IVS-mCD8::GFP* cut with *NotI/XbaI*. The plasmid was integrated into the *D. melanogaster* genome at the *attP2* site through phiC31-mediated transgenesis by BestGene Inc.

### Calcium imaging

We used the calcium imaging procedure described in^52^ with several modifications. In brief, female flies aged 5-9 days after eclosion were mounted and their head cuticle, tracheae, and glands above the brain were removed in adult hemolymph (AHL) saline (108 mM NaCl, 5 mM KCl, 2 mM CaCl_2_, 8.2 mM MgCl_2_, 4 mM NaHCO_3_, 1 mM NaH_2_PO_4_, 5 mM trehalose, 10 mM sucrose, 5 mM HEPES, pH 7.5). Odors were diluted in dichloromethane solvent to 1% (v/v). Images were acquired using a commercial upright microscope (Zeiss LSM 710 NLO) either using the two-photon or one-photon mode. An upright Zeiss AxioExaminer Z1 was fitted with a Ti:Sapphire Chameleon Ultra II infra-red laser (Coherent) for the two-photon mode (all experiments except for *Cngl^RNAi^* and Figure S3F) and 488 nm laser for the one-photon mode (*Cngl^RNAi^* and Figure S3F) as excitation source. Images were acquired with a 20× water dipping objective (Plan-Apochromat 20× W; NA 1.0), with a resolution of 128 × 128 pixels (0.8926 pixels/µm) and a scan speed of 6.30 µs/pixel^2^. For the two-photon mode, the excitation wavelength was set to 930 nm. Each measurement consisted of 50 images acquired at 4.17 Hz, with stimulation starting ∼5 s after the beginning of the acquisition and lasting for 1 s. Antennae were stimulated using a custom-made olfactometer. For low airflow experiments (i.e., all experiments if not stated otherwise), antennae were permanently exposed to air at a flow rate of 0.48 l/min by combining a main airstream that carries the odor (0.24 l/min) and a secondary stream (0.24 l/min) of room air. For high airflow experiments (i.e., **Extended Data** Figure 1b, **Extended Data** Figure 1d and **Extended Data** Figure 4a), antennae were permanently exposed to air at a flow rate of 1.5 l/min by combining a main airstream carrying the odor (0.5 l/min) and a secondary stream (1 l/min) of room air.

Data were processed using Fiji and custom-written scripts in R. The signal intensity averaged across the ROI (ovals contained in each glomerulus; example in **Extended Data** Figure 1a) for each timeframe (hereafter *F*) was used to calculate the normalized signal *ΔF/F_0_* . Here, *F_0_* (baseline fluorescence) was calculated as the average *F* during frames 16–19 (1 s before olfactory stimulus onset). The peak *ΔF/F_0_* value (which represents the odor response intensity) was calculated as the maximum *ΔF/F_0_* value during frames 20–29 (2.5 s after olfactory stimulation onset). Then, the *ΔF/F_0_* values were normalized for each animal by the maximum *ΔF/F_0_* value obtained across different odors to correct for varying basal GCaMP fluorescence across genotypes. Principal component analyses (PCA) were performed based on the odor responses (maximum *ΔF/F_0_* values) after subtracting the solvent response for each neuron type in each animal. The Euclidean distance was calculated by squaring the odor response difference between Ir64a-DC4 and Ir64a-DP1m for each odor, summing them across odors, and taking a square root in each animal.

Odors used are listed in **Extended Data Table 4.**

### Immunohistochemistry

Antennae were collected by flash-freezing the flies in a mini-sieve (#378460000, bel-art products) with liquid nitrogen and tapping the sieve on top of a Petri dish filled with fixative solution (PBS + 4% PFA, 3% Triton-X-100). The antennae were transferred into a 2 ml microcentrifuge tube and fixed for 2-3 h at 4°C with rotation. The antennae were then washed briefly 3× in PBS + 3% Triton-X-100 and once in PBS + 0.1% Triton-X-100 (PTX) at room temperature (RT). Following 3× 10 min washes with PTX at RT, antennae were blocked for 1 h in PTX + 5% heat inactivated normal Goat serum at RT. The blocking solution was then replaced with primary antibody diluted in blocking solution and incubated for 3 days at 4°C. After washing 6× 10 min with PTX at RT, blocking was performed as previously and then replaced with secondary antibody diluted in blocking solution and incubated for 2 days at 4°C. After washing 6× 10 min with PTX at RT, mounting medium (Vectashield Antifade Mounting Medium (H-1000-10), Vector Laboratories) was added and allowed to permeate tissue for 1 day. Antennae were mounted on a glass slide with a bridge made of nail polish to prevent tissue being crushed by the cover slip.

Brains were dissected in PB and fixed with PB + 4% PFA for 25 min at RT in a glass dissection well, prior to transfer to a 1.5 ml microcentrifuge tube and 6 × 15 min washes in PB + 0.1% Triton-X-100 (PBT) at RT. The brains were then blocked for 1 h in PBT + 5% heat inactivated normal goat serum at RT. The blocking solution was replaced with primary antibody diluted in blocking solution and incubated for 3 days at 4°C. After washing 6× 15 min with PBT at RT, the solution was replaced with secondary antibody diluted in blocking solution and incubated for 2 days at 4°C. After washing 6× 15 min with PBT at RT, mounting medium was added and allowed to permeate tissue for 1 day. Brains were then mounted on a glass slide with a bridge of cover slips.

Antibodies used are listed in **Extended Data Table 5**.

### Image acquisition and processing

Confocal images of antennae and brains were acquired on an inverted confocal microscope (Zeiss LSM 880 Airyscan) with an oil immersion 63× objective (Plan Apochromat 63× 1.4 Oil DIC). 0.7/0.8× and 3× zoom images were taken for whole antenna and sacculus, respectively. 0.6× and 0.8× zoom images were taken for imaging of central brains and antennal lobe, respectively. For the images used for signal quantification, the laser power and gain setting were kept consistent for all samples across genotypes.

The quantification of fluorescence signals from the images were performed using the Fiji software and the analyses were performed blindly to the genotype through file name encryption. The sensilla of sacculus chamber III were identified as GS1 (Ir64a-DC4) or GS2 (Ir64a-DP1m) based on their location and morphology from the transmitted light channel: sacculus chamber III is separated by a cuticular flap and houses thick, blunter GS1 resides in the proximal compartment and slender, sharper GS2 in the distal compartment (**Figure 1a**)^5,17^. The ROIs were set as rectangles that fit inside each sensilla. Signals from more than three sensilla were averaged for each type and for each antenna.

### snRNA-seq data

Most analyses used processed data from the control developmental antennal snRNA-seq atlas^19^, which were analyzed in Seurat (v5.3.0). With the exception of **Extended Data** Figure 3d, analysis of gene expression levels used data only from the “late” developmental phase, using the subset function implemented in Seurat.

The comparative antennal atlases of *D. melanogaster* (Canton-S) and *D. sechellia* (DSSC 14021-0248.07) will be described in more detail in a separate paper. In brief, three replicates of 100-200 third antennal segments were harvested by snap-freezing animals in a mini-sieve with liquid nitrogen and agitating the animals to break off the appendages. Single-nucleus suspensions were prepared and sorted following the Fly Cell Atlas protocol^33^ and 15,000–20,000 nuclei were loaded onto a 10x Genomics Chromium Next GEM Chip to target 10,000 nuclei for sequencing. Single-nucleus RNA-seq libraries were generated using the Chromium Single Cell 3’ v3.1 dual-index kit. Libraries were pooled for cluster generation on the Illumina NovaSeq 10x platform and sequenced according to 10x Genomics guidelines. Demultiplexing was performed using bcl2fastq2 v2.20. DEGs were identified using the FindMarkers function in Seurat (test.use = “MAST”, min.pct = 0.05). Species datasets were integrated and clustered using Seurat. The Ir-expressing cluster, defined by the Ir co-receptor marker genes *Ir25a*, *Ir8a*, and *Ir76b*, was subsequently subclustered, yielding multiple Ir-expressing populations, including two distinct *Ir64a*-positive neuronal subtypes.

### Quantification and statistical analysis

The statistical analyses used are indicated in respective figure legends. The statistical tests were performed using R/RStudio.

---

**Extended Data Figure 1.**
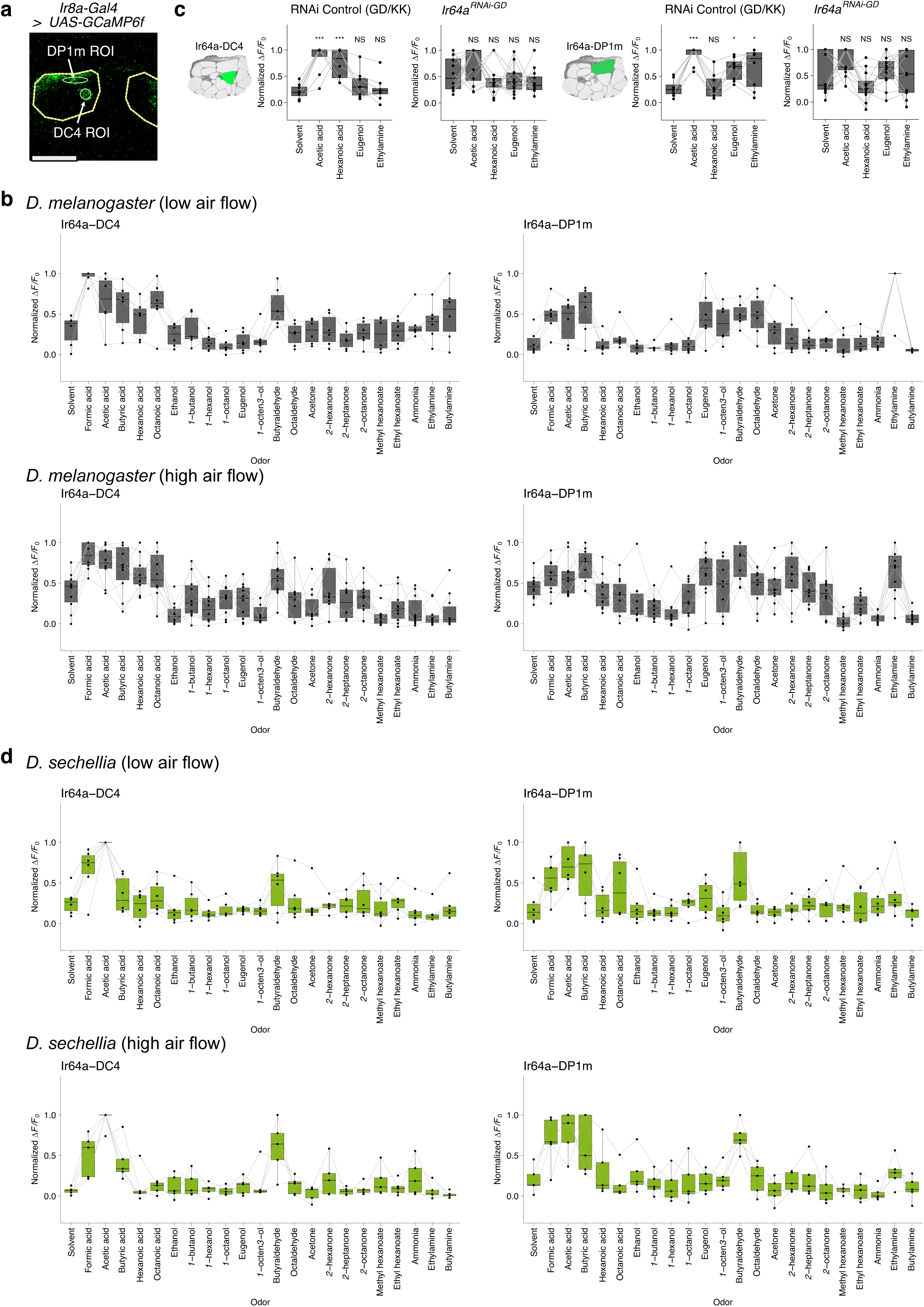
Odor tuning profiles of Ir64a neurons in *D. melanogaster* and *D. sechellia*. a Representative regions of interest (ROIs) used to quantify GCaMP signals from Ir64a-DC4 and Ir64a-DP1m neurons. Dotted yellow line marks the outline of the antennal lobe. Scale bar = 50 µm. Genotype: *w;Ir8a-Gal4/UAS-GCaMP6f*. **b** Quantifications of calcium responses to the indicated odors in *D. melanogaster* at low (0.24 l/min) and high (1.0 l/min) air flow, with the latter likely increasing odor concentration arriving at the fly and thus enhancing responses. *n* = 6 (low air flow) and 10 (high air flow) animals. Genotype: *w;Ir8a-Gal4/UAS-GCaMP6f* **c** Quantifications of calcium responses to the indicated odors in *Ir64a^RNAi^* animals. *n* = 9 animals. The control data are replotted from Figure 1f as a reference. Genotype: *w/w,UAS-Dcr-2;Ir64a-Gal4,UAS-GCaMP6f/+;UAS-Ir64a^RNAi^/+*. **d** Quantifications of calcium responses to the indicated odors in *D. sechellia* at low (0.24 l/min) and high (1.0 l/min) air flow. *n* = 6 (low air flow) and 5 (high air flow) animals. Genotype: *w/w,UAS-GCaMP6f;Ir8a-Gal4/+* (chromosome unidentified).

**Extended Data Figure 2.**
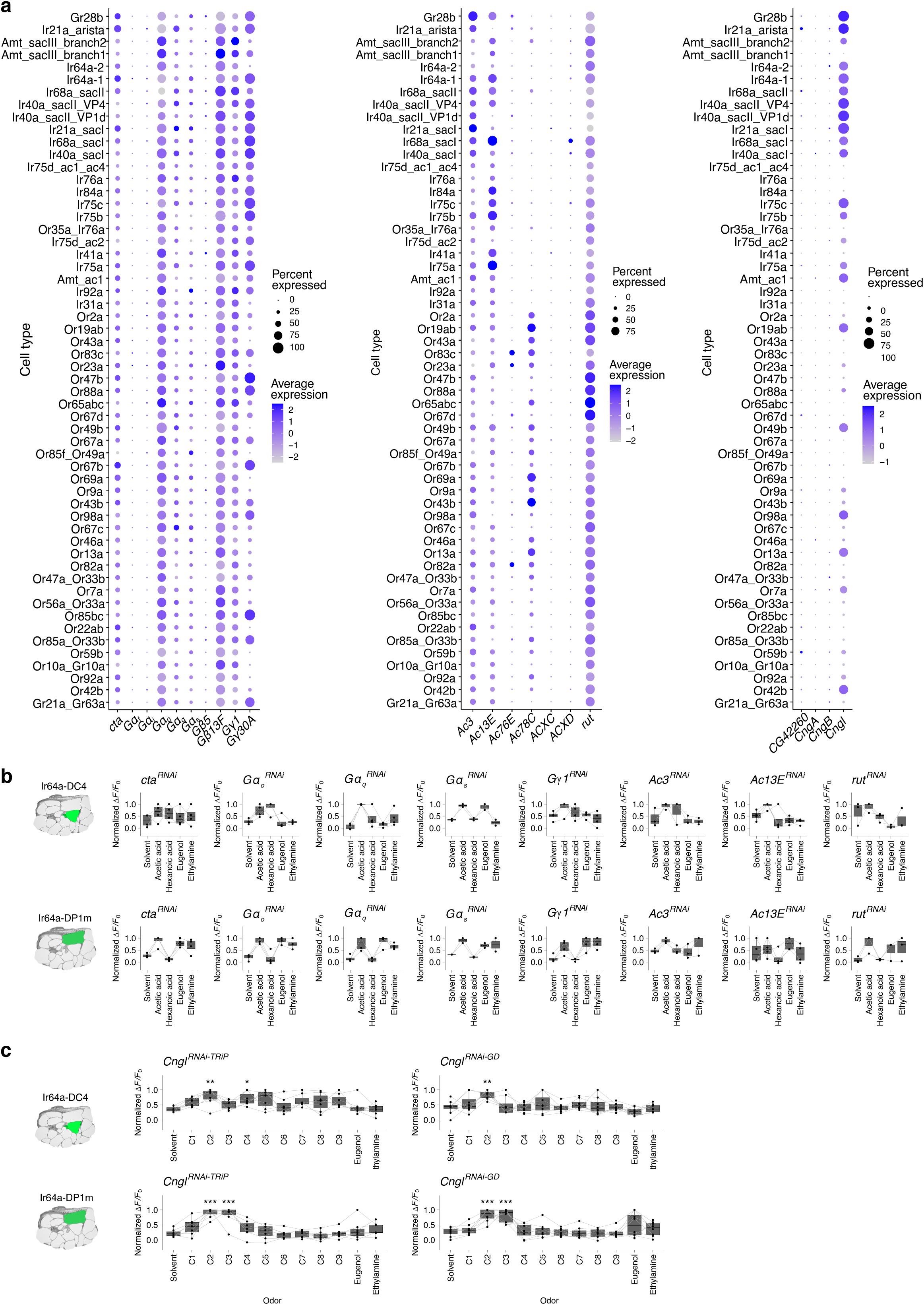
Expression and function of metabotropic signaling molecules in in sensory neurons. a Dot plots showing the expression of genes encoding G protein subunits, adenylate cyclases and cyclic nucleotide-gated ion channels in OSNs. Genes undetectable in all populations were excluded. Broad expression of several G protein subunits is concordant with previous immunohistochemical analyses^54^. **b** Quantifications of calcium responses to the indicated odors upon knock-down of the indicated G protein subunits and adenylate cyclases. Small-scale screening results (i.e., with limited *n*) are shown. Genotypes: *y,v/w,UAS-Dcr-2;Ir64a-Gal4,UAS-GCaMP6f/UAS-cta^RNAi^* (*cta^RNAi^*) | *y,sc,v,sev/w,UAS-Dcr-2;Ir64a-Gal4,UAS-GCaMP6f/+;UAS-Gα_o_^RNAi^/+* (*Gα_o_^RNAi^*) | *y,v/w,UAS-Dcr-2;Ir64a-Gal4,UAS-GCaMP6f/+;UAS-Gα_q_^RNAi^/+* (*Gα_q_^RNAi^*) | *y,v/w,UAS-Dcr-2;Ir64a-Gal4,UAS-GCaMP6f/+;UAS-Gα_s_^RNAi^/+* (*Gα_s_^RNAi^*) | *y,sc,v,sev/w,UAS-Dcr-2;Ir64a-Gal4,UAS-GCaMP6f/+;UAS-Gγ1^RNAi^/+* (*Gγ1^RNAi^*) | *y,v/w,UAS-Dcr-2;Ir64a-Gal4,UAS-GCaMP6f/UAS-Ac3^RNAi^*(*Ac3^RNAi^*) | *y,sc,v,sev/w,UAS-Dcr-2;Ir64a-Gal4,UAS-GCaMP6f/UAS-Ac13E^RNAi^* (*Ac13E^RNAi^*) | *y,v/w,UAS-Dcr-2;Ir64a-Gal4,UAS-GCaMP6f/UAS-rut^RNAi^* (*rut^RNAi^*). **c** Quantifications of calcium responses to the indicated odors upon *Cngl* knock-down. *n* = 7 (*Cngl^RNAi-TRiP^*) and 8 (*Cngl^RNAi-GD^*) animals. Genotypes: *y,v/w,UAS-Dcr-2;Ir64a-Gal4,UAS-GCaMP6f/+;UAS-Cngl^RNAi-TRiP^/+* (*Cngl^RNAi-TRiP^*) | *w/w,UAS-Dcr-2;Ir64a-Gal4,UAS-GCaMP6f/UAS-Cngl^RNAi-GD^* (*Cngl^RNAi-GD^*). Dunnett’s test (two-sided, control: solvent), \*\*\**P* < 0.001, \*\**P* < 0.01, \**P* < 0.05, otherwise *P* > 0.05.

**Extended Data Figure 3.**
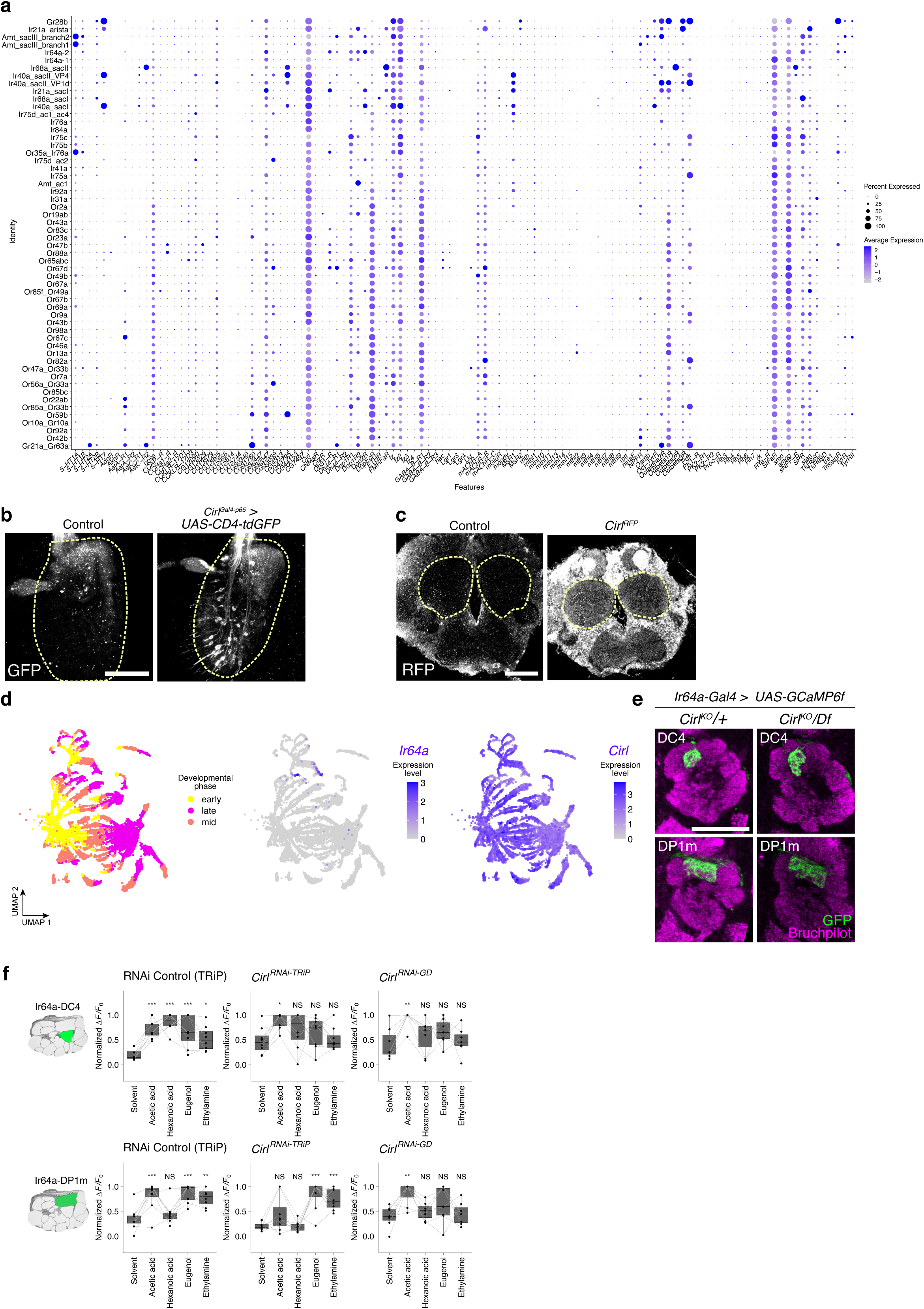
Characterization of Cirl expression and function. **a** Dot plot showing the expression of *D. melanogaster* GPCR genes in all OSN types. The GPCR gene list was obtained from FlyBase^55^ (https://flybase.org/reports/FBgg0000172). **b** Expression pattern in the antenna of CD4:tdGFP driven by *Cirl^Gal4-p65^* (visualized with anti-GFP). Scale bar = 50 µm. Genotypes: *yw/w;UAS-CD4:tdGFP/+* (control) | *yw/w;UAS-CD4:tdGFP/Cirl^Gal4-p65^*. The Gal4 driver is active in many, but far from all, OSNs, at least partially concordant with the broad *Cirl* transcript expression (Figure 3b). **c** Representative image of Cirl^RFP^ localization (visualized with anti-RFP) in the brain. Dotted yellow line marks the outline of the antennal lobe. Scale bar = 50 µm. Genotypes: *w^1118^* (control)*| w^1118^*;*Cirl^RFP.V5^*(*Cirl^RFP^*). **d** Left: UMAP of all sensory neuron lineages in the antennal developmental atlas, color-coded by developmental phase^19^. Right: the same UMAP illustrating expression of *Ir64a* and *Cirl*. **e** Representative images of Ir64a glomeruli (DC4 and DP1m) in *Cirl* heterozygous (*Cirl^KO^/+*) and hemizygous (*Cirl^KO^/Df*) mutants visualized with anti-GFP (detecting GCaMP6f) and anti-Bruchpilot (nc82). Scale bar = 50 µm. Genotypes: *w;UAS-GCaMP6f,Cirl^KO^/+;Ir64a-Gal4/+* | *w;UAS-GCaMP6f,Cirl^KO^/Df(2R)Exel8047;Ir64a-Gal4/+*. Here, GCaMP6f is simply used as a visual marker of the axon termini of Ir64a neurons. **f** Quantifications of calcium responses to the indicated odors upon knock-down of *Cirl*. *n* = 8 (control), 8 (*Cirl^RNAi-TRiP^*), 7 (*Cirl^RNAi-GD^*), 7 (*Cirl^RNAi-KK^*) animals. Genotypes: *y,sc,v,sev/w,UAS-Dcr-2;Ir64a-Gal4,UAS-GCaMP6f/+;UAS-mCherry.VALIUM10/+* (control) | *y,v/w,UAS-Dcr-2;Ir64a-Gal4,UAS-GCaMP6f/+;UAS-Cirl^RNAi-TRiP^/+* (*Cirl^RNAi-TRiP^*) | *w/w,UAS-Dcr-2; Ir64a-Gal4,UAS-GCaMP6f/UAS-Cirl^RNAi-GD^* (*Cirl^RNAi-^*^GD^).

**Extended Data Figure 4.**
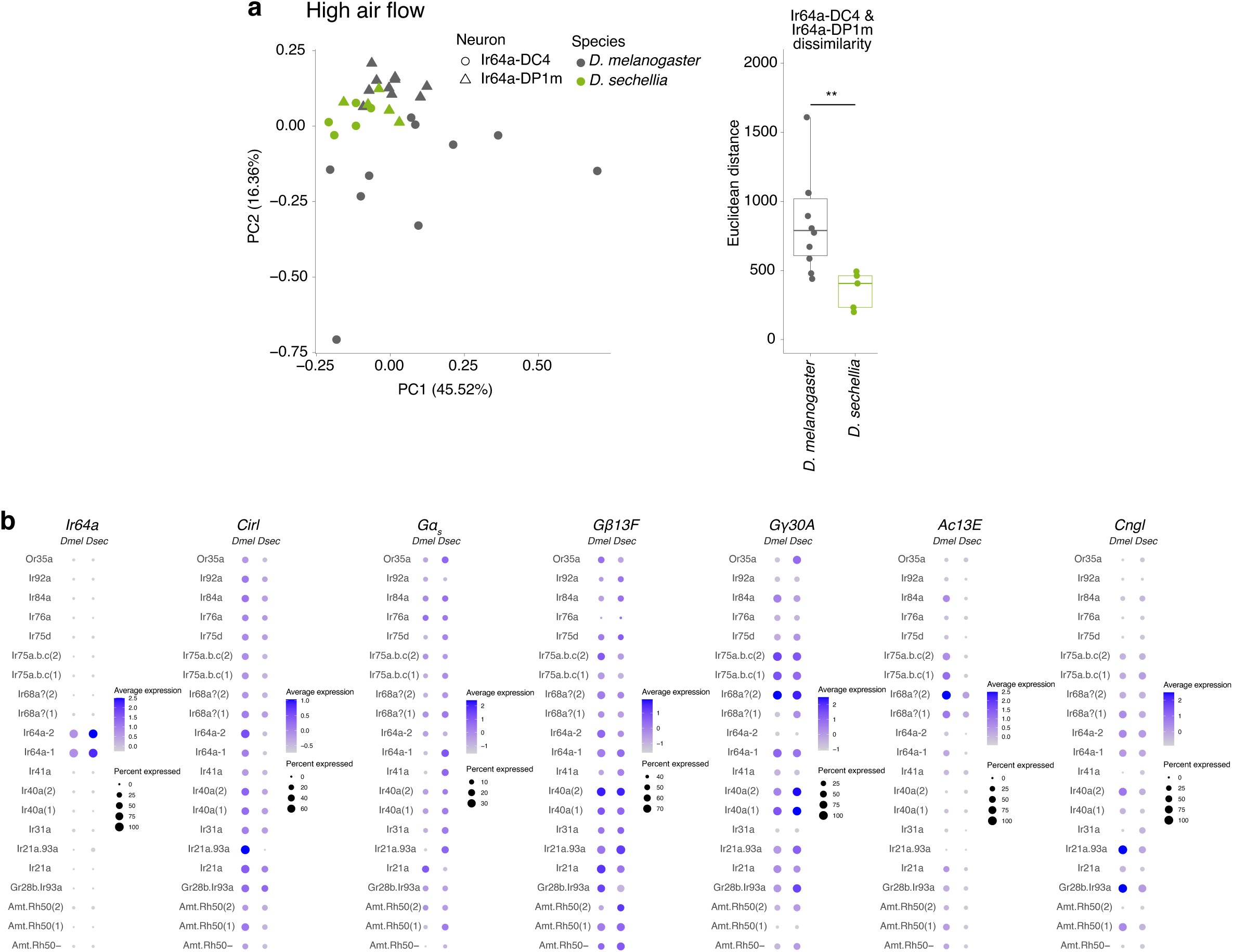
Interspecific analysis of functional and transcriptional divergence in Ir64a neuron subtypes. a Left: Principal component analysis of odor tuning profile based on broader odor responses (**Extended Data** Figure 1b,d) in both Ir64a neuron subtypes and both drosophilid species at high air flow (1 l/min). Right: quantifications of the Euclidean distance between Ir64a-DC4 and Ir64a-DP1m neuron responses in each species. **b** Dot plot showing the expression of the indicated genes in various antennal neuron populations of the Ir subsystem of *D. melanogaster* and *D. sechellia*.

**Extended Data Table 1.** Differentially-expressed genes between *D. melanogaster* Ir64a neuron subtypes. - see separate Excel document

**Extended Data Table 2.** Differentially-expressed genes between *D. melanogaster* and *D. sechellia* Ir64a-1 neurons. - see separate Excel document

**Extended Data Table 3.**
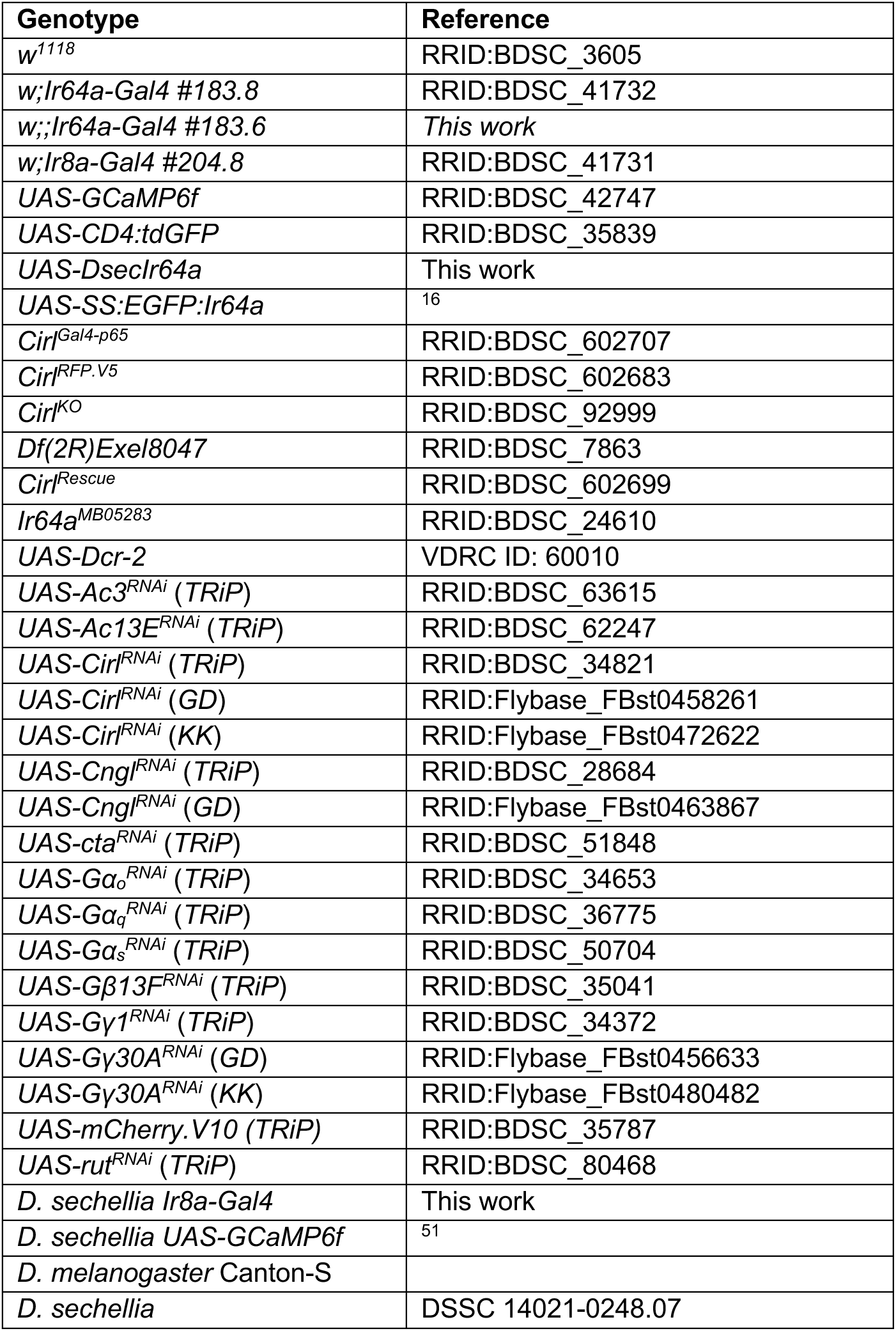
*Drosophila* strains.

| Genotype | Reference |
| --- | --- |
| *w*<sup>1118</sup> | RRID:BDSC_3605 |
| *w;Ir64a-Gal4* #183.8 | RRID:BDSC_41732 |
| *w;;Ir64a-Gal4* #183.6 | *This work* |
| *w;Ir8a-Gal4* #204.8 | RRID:BDSC_41731 |
| *UAS-GCaMP6f* | RRID:BDSC_42747 |
| *UAS-CD4:tdGFP* | RRID:BDSC_35839 |
| *UAS-DsecIr64a* | *This work* |
| *UAS-SS:EGFP:Ir64a* | <sup>16</sup> |
| *CirI*<sup>Gal4-p65</sup> | RRID:BDSC_602707 |
| *CirI*<sup>RFP.V5</sup> | RRID:BDSC_602683 |
| *CirI*<sup>KO</sup> | RRID:BDSC_92999 |
| *Df(2R)Exel8047* | RRID:BDSC_7863 |
| *CirI*<sup>Rescue</sup> | RRID:BDSC_602699 |
| *Ir64a*<sup>MB05283</sup> | RRID:BDSC_24610 |
| *UAS-Dcr-2* | VDRC ID: 60010 |
| *UAS-Ac3<sup>RNAi</sup> (TRiP)* | RRID:BDSC_63615 |
| *UAS-Ac13<sup>E<sup>RNAi</sup></sup> (TRiP)* | RRID:BDSC_62247 |
| *UAS-CirI<sup>RNAi</sup> (TRiP)* | RRID:BDSC_34821 |
| *UAS-CirI<sup>RNAi</sup> (GD)* | RRID:Flybase_FBst0458261 |
| *UAS-CirI<sup>RNAi</sup> (KK)* | RRID:Flybase_FBst0472622 |
| *UAS-Cngl<sup>RNAi</sup> (TRiP)* | RRID:BDSC_28684 |
| *UAS-Cngl<sup>RNAi</sup> (GD)* | RRID:Flybase_FBst0463867 |
| *UAS-cta<sup>RNAi</sup> (TRiP)* | RRID:BDSC_51848 |
| *UAS-Ga<sub>o</sub><sup>RNAi</sup> (TRiP)* | RRID:BDSC_34653 |
| *UAS-Ga<sub>q</sub><sup>RNAi</sup> (TRiP)* | RRID:BDSC_36775 |
| *UAS-Ga<sub>s</sub><sup>RNAi</sup> (TRiP)* | RRID:BDSC_50704 |
| *UAS-Gβ13<sup>F<sup>RNAi</sup></sup> (TRiP)* | RRID:BDSC_35041 |
| *UAS-Gγ1<sup>RNAi</sup> (TRiP)* | RRID:BDSC_34372 |
| *UAS-Gγ30A<sup>RNAi</sup> (GD)* | RRID:Flybase_FBst0456633 |
| *UAS-Gγ30A<sup>RNAi</sup> (KK)* | RRID:Flybase_FBst0480482 |
| *UAS-mCherry.V10 (TRiP)* | RRID:BDSC_35787 |
| *UAS-rut<sup>RNAi</sup> (TRiP)* | RRID:BDSC_80468 |
| *D. sechellia Ir8a-Gal4* | *This work* |
| *D. sechellia UAS-GCaMP6f* | <sup>51</sup> |
| *D. melanogaster* Canton-S |  |
| *D. sechellia* | DSSC 14021-0248.07 |

**Extended Data Table 4.** Odors.

| Odor | Reference/source | CAS |
| --- | --- | --- |
| Dichloromethane (solvent) | Thermo scientific 406920010 | 75-09-2 |
| Formic acid | SIGMA-ALDRICH 33015 | 64-18-6 |
| Acetic acid | FLUKA 45726 | 64-19-7 |
| Propionic acid | SIGMA-ALDRICH P1386 | 79-09-4 |
| Butyric acid | SIGMA-ALDRICH B103500 | 107-92-6 |
| Valeric acid | SIGMA-ALDRICH 240370 | 109-52-4 |
| Hexanoic acid | SIGMA-ALDRICH 153745 | 142-62-1 |
| Heptanoic acid | FLUKA 75190 | 111-14-8 |
| Octanoic acid | SIGMA-ALDRICH C2875 | 124-07-2 |
| Nonanoic acid | FLUKA 76342 | 112-05-0 |
| Ethanol | CARLO ERBA 4146082 | 64-17-5 |
| 1-butanol | SIGMA-ALDRICH 537993 | 71-36-3 |
| 1-hexanol | SIGMA-ALDRICH 471402 | 111-27-3 |
| 1-octanol | FLUKA 74848 | 111-87-5 |
| Eugenol | FLUKA 46100 | 97-53-0 |
| 1-octen-3-ol | FLUKA 74950 | 3391-86-4 |
| Butyraldehyde | FLUKA 20711 | 123-72-8 |
| Octaldehyde | SIGMA-ALDRICH O5608 | 124-13-0 |
| Acetone | SIGMA-ALDRICH 179124 | 67-64-1 |
| 2-hexanone | SIGMA-ALDRICH 103004 | 591-78-6 |
| 2-heptanone | SIGMA-ALDRICH 537683 | 110-43-0 |
| 2-octanone | SIGMA-ALDRICH O4709 | 111-13-7 |
| Methyl hexanoate | SIGMA-ALDRICH 259942 | 106-70-7 |
| Ethyl hexanoate | SIGMA-ALDRICH 148962 | 123-66-0 |
| Ammonia | SIGMA-ALDRICH 320145 | 1336-21-6 |
| Ethylamine | FLUKA 02950 | 75-04-7 |
| Butylamine | SIGMA-ALDRICH 471305 | 109-73-9 |

**Extended Data Table 5.** Antibodies.

| Antibody | Dilution | Reference/source | Identifier |
| --- | --- | --- | --- |
| Rabbit anti-RFP | 1:250 | Abcam | ab62341 |
| Rabbit anti-Ir64a | 1:500 | <sup>6</sup> | RRID:AB_2566854 |
| Rabbit anti-GFP | 1:500 | Invitrogen | A-6455 |
| Mouse anti-GFP | 1:500 | Invitrogen | A11120 |
| Mouse anti-Bruchpilot (nc82) | 1:10 | DSHB | RRID:AB_2392664 |
| Alexa Fluor 488 Goat anti-Rabbit | 1:500 | Invitrogen | A11034 |
| Alexa Fluor 488 Goat anti-Mouse | 1:500 | Invitrogen | A11029 |
| Cy5 Goat anti-Rabbit | 1:500 | Jackson ImmunoResearch | 111-175-144 |
| Cy5 Goat anti-Mouse | 1:500 | Jackson ImmunoResearch | 115-175-166 |

